# Functional interaction of torsinA and its activators in liver lipid metabolism

**DOI:** 10.1101/2023.06.21.545957

**Authors:** Antonio Hernandez-Ono, Yi Peng Zhao, John W. Murray, Cecilia Östlund, Michael J. Lee, Angsi Shi, William T. Dauer, Howard J. Worman, Henry N. Ginsberg, Ji-Yeon Shin

## Abstract

TorsinA is an atypical ATPase that lacks intrinsic activity unless it is bound to its activators lamina-associated polypeptide 1 (LAP1) in the perinuclear space or luminal domain-like LAP1 (LULL1) throughout the endoplasmic reticulum. However, the interaction of torsinA with LAP1 and LULL1 has not yet been shown to modulate a defined physiological process in mammals *in vivo*. We previously demonstrated that depletion of torsinA from mouse hepatocytes leads to reduced liver triglyceride secretion and marked steatosis, whereas depletion of LAP1 had more modest similar effects. We now show that depletion of LULL1 alone does not significantly decrease liver triglyceride secretion or cause steatosis. However, simultaneous depletion of both LAP1 and LULL1 from hepatocytes leads to defective triglyceride secretion and marked steatosis similar to that observed with depletion of torsinA. Our results demonstrate that torsinA and its activators dynamically regulate a physiological process in mammals *in vivo*.

## Introduction

TorsinA is an ATPase associated with various cellular activities (AAA+) that resides in the endoplasmic reticulum (ER) lumen and continuous perinuclear space of the nuclear envelope ^1–3^. TorsinA interacts with at least two differentially localized transmembrane proteins that contain highly conserved luminal domains. In the perinuclear space, it interacts with lamina-associated polypeptide 1 (LAP1) at the inner nuclear membrane and with luminal domain-like LAP1 (LULL1) throughout the entire ER ^4^. *In vitro*, torsinA does not display ATPase activity in isolation; however, ATP hydrolysis is induced upon association with the luminal domains of LAP1 or LULL1 ^5, 6^. Structural biology studies have shown that torsinA combines with these transmembrane proteins to form heterohexameric rings in which LAP1 and LULL1 induce ATPase activity by supplying an arginine to the neighboring torsinA molecule ^6^. A 3-basepair deletion (ΔGAG) in *TOR1A* encoding torsinA, resulting in loss of a glutamic acid at residues 302/303 in the carboxyl-terminal of torsinA, causes the autosomal dominant movement disorder early-onset DYT1 dystonia ^7^. Data from genetically-modified mice demonstrate that this variant leads to loss of torsinA function in maintaining neuronal nuclear envelope morphology ^8^. The role of LAP1 and LULL1 in activating torsinA is further supported by the finding that deletion of glutamic acid 302/303 from torsinA impairs its ATP-dependent interaction with these proteins ^9, 10^. However, even with the knowledge provided by these elegant biochemical, structural biological, and cell biology studies, the *in vivo* functional relevance of the interaction between torsinA and its activators is largely unknown.

We have identified novel roles for torsinA and LAP1 in liver lipid metabolism. Mice with hepatocyte-specific depletion of torsinA (A-CKO mice) have reduced secretion of triglyceride (TG)-rich very low-density lipoprotein (VLDL) and marked steatosis, while livers from mice with hepatocyte-specific LAP1 deletion (L-CKO mice) demonstrate more modest reductions in VLDL secretion and less severe steatosis ^11^. We took advantage of these mammalian systems with measurable readouts to investigate the unresolved functional significance of the torsinA activators LAP1 and LULL1 *in vivo*. Given the more modest decrease in VLDL secretion and less severe steatosis in livers of L-CKO mice compared to A-CKO mice, we reasoned that LULL1 may provide some degree of torsinA activation in the absence of LAP1. We therefore hypothesized that the combined depletion of LAP1 and LULL1 from hepatocytes is sufficient to abolish all torsinA function and phenocopy A-CKO mice that have depletion of torsinA from hepatocytes. We now test this hypothesis *in vivo*.

## Results

### Mice with depletion of LULL1 from hepatocytes have no steatosis and normal VLDL secretion

A-CKO mice with hepatocyte-specific depletion of torsinA develop striking hepatic steatosis, while L-CKO mice with depletion of LAP1 have a relatively mild steatosis phenotype ^11^. This suggests that LULL1 may activate torsinA in the ER or continuous nuclear envelope, providing partial cellular activity. To investigate the potential role of LULL1 in hepatic steatosis and lipoprotein secretion, we crossed mice with two floxed alleles of *Tor1aip2* encoding LULL1 (LUL-flox mice) to mice with an albumin-Cre transgene to obtain mice with depletion of LULL1 from hepatocytes (LUL-CKO mice) (Supplementary Fig. 1). Both male and female LUL-CKO mice developed normally without overt abnormal phenotypes. Given the absence of apparent sex-specific differences, we subsequently analyzed them in an aggregated manner. Livers from LUL-CKO mice at 4 months of age were grossly normal (Fig. 1a). Immunoblotting showed that LULL1 expression was undetectable in livers from these mice (Fig. 1b). In control mice, our anti-LULL1 antibody detected two closely migrating bands as shown in a previous report ^8^. On histological examination, liver sections from LUL-CKO mice stained with hematoxylin and eosin or Oil Red O to detect fat did not appear different than those from littermate controls (Fig. 1c). Based on interpretation of the sections stained with hematoxylin and eosin, a pathologist blind to genotype scored liver steatosis (on a scale of 0 to 3) as 0.5 ± 0.14 in the LUL-flox mice and 0.75 ± 0.13 in LUL-CKO mice (mean ± SEM, n = 4, *p*-value not significant by Student’s t-test). The steady state liver TG content in LUL-CKO mice was also similar to that in control mice (Fig. 1d). To assess liver VLDL secretion, we measured plasma TG concentrations following intravenous administration of tyloxapol to block both lipolysis of VLDL TG and uptake of circulating TG-rich VLDL remnant lipoproteins. There were no differences in plasma TG concentrations between LUL-CKO and controls at all time points measured after tyloxapol administration (Fig. 1e). The calculated hepatic TG secretion rate was the same in LUL-CKO and control mice (Fig. 1f). Analysis of plasma proteins by SDS-PAGE followed by autoradiography 120 minutes after injection of ^35^S-methionine with tyloxapol showed no significant differences of radiolabeled apolipoprotein (apo) B100 or apoB48 (Fig. 1g and 1h). ApoB100 and apoB48 are major protein components of mouse VLDL ^12, 13^. These results show that depletion of LULL1 from hepatocytes does not lead to steatosis or decreased VLDL secretion.

**Fig. 1.**
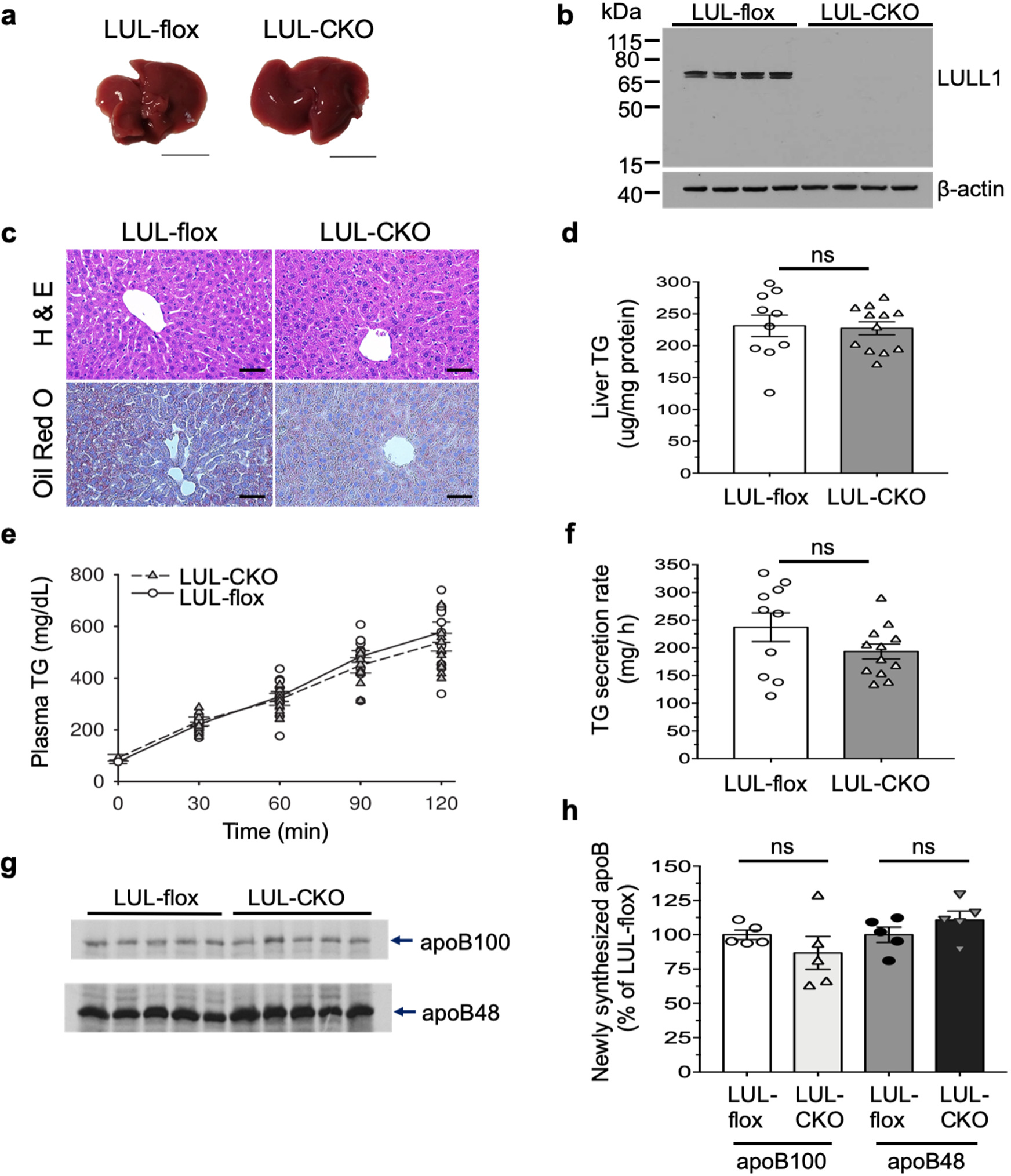
Mice with depletion of LULL1 from hepatocytes show no overt liver abnormalities. **a** Representative photographs of livers from 4-month-old control (LUL-flox) and LUL-CKO mice fed a chow diet. Bars: 1 cm. **b** Immunoblots of liver lysates from LUL-flox (control) and LUL-CKO mice probed with antibodies against LULL1 and β-actin. The anti-LULL1 antibody detects two closely migrating bands ^8^. **c** Representative light photomicrographs of liver sections from chow fed mice stained with hematoxylin and eosin (H & E) or Oil Red O. Scale bars: 50 μm. **d** Liver TG content of LUL-flox (n = 10) and LUL-CKO (n = 12) mice. Mice were fasted for 4-5 hours before collecting livers to measure TG content. **e** Plasma TG concentration versus time after injection of tyloxapol to block peripheral uptake in LUL-CKO (n = 12) and LUL-flox (n = 10) mice. **f** TG secretion rates calculated from the changes in plasma concentrations between 30 and 120 minutes in e. **g** Autoradiogram of SDS-polyacrylamide gel showing ^35^S-labeled plasma proteins collected 120 minutes after injection with ^35^S-methionine and tyloxapol. Each lane shows proteins from an individual mouse. Migrations of ^35^S-methionine-labeled apoB100 and apoB48 are indicated (n = 5 mice per group). **h** Bands corresponding to apoB100 and apoB48 shown in f were quantified by densitometry and shown as a % of the value from LUL-flox group (n = 5 mice per group). In panels d, e, f and h, values are means ± SEM with each circle or triangle representing the value from an individual mouse. None of the differences were statistically significant (ns) by Student’s t-test.

### Hepatocytes from LUL-CKO mice do not have increased TG accumulation

LULL1 overexpression promotes torsinA redistribution to the nuclear envelope in cultured cells ^10, 14^. Therefore, we examined if the expression or distribution of endogenous torsinA or LAP1 is changed in LULL1-deficient hepatocytes. Immunoblotting of whole liver lysates from LUL-CKO mice showed depletion of LULL1, no changes in LAP1, and a slight but significant increase in torsinA expression (Supplementary Fig. 2a and 2b). Immunoblotting of lysates of primary hepatocytes isolated from LUL-CKO mice showed depletion of LULL1, a slight decrease in LAP1, and a slight but significant increase in torsinA expression (Fig. 2a and 2b). Immunofluorescence microscopy did not identify obvious changes in the distribution of endogenous LAP1 or torsinA in hepatocytes isolated from LUL-CKO mice (Fig. 2c). BODIPY staining of neutral lipids in hepatocytes isolated from LUL-CKO mice did not show nuclear lipid droplets, which are observed in mice with hepatocyte-specific depletion of LAP1 ^11, 15^, or an increase of cytosolic lipid content compared to control (Fig. 2d). We quantified BODIPY fluorescence intensity using an automated measuring method (Supplementary Fig. 2c). The results did not show a difference in mean fluorescence intensities in LUL-flox and LUL-CKO mice (Supplementary Fig. 2d). These results show that depletion of LULL1 from hepatocytes does not lead to excess lipid accumulation and that LAP1 may be sufficient to stimulate torsinA activity when LULL1 is absent.

**Fig. 2.**
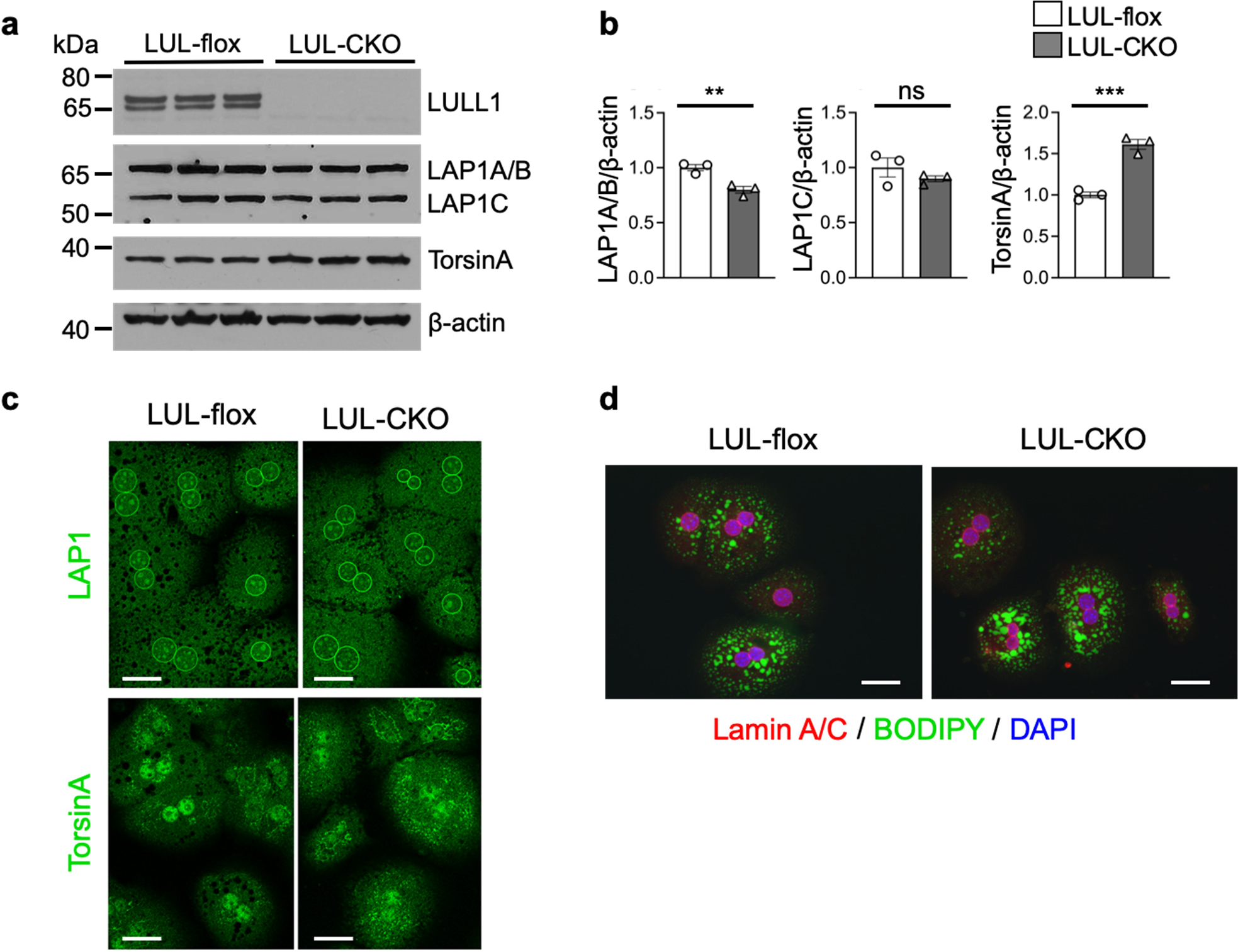
Expression of torsinA and LAP1 in livers and hepatocytes of LUL-CKO mice. **a** Immunoblots of cell lysates from hepatocyte cultures from LUL-flox and LUL-CKO mice at 4 months of age. Blots were probed with antibodies against LULL1, LAP1, torsinA, and β-actin. **b** Relative ratios of band densities of LAP1A/B, LAP1C, or torsinA to β-actin blots shown in panel c (n = 3 different hepatocyte cultures from 1 mouse each group). Values are means ± SEM with individual circle and triangle representing the value from each lane on the immunoblots in panel a. \*\**p* < 0.01, \*\*\**p* < 0.001, ns = not significant by Student’s t-test. **c** Confocal immunofluorescence micrographs of isolated hepatocytes labeled with anti-LAP1 (green, upper panel) and anti-torsinA (green, bottom panel) antibodies. Scale bars: 20 μm. **d** Confocal immunofluorescence micrographs of hepatocytes stained with anti-lamin A/C antibody (red), BODIPY (green) and DAPI (blue). Scale bars: 20 μm.

As hepatocytes isolated from LUL-CKO mice did not differ in lipid content compared to LUL-flox controls, we tested if this was a result of lower expression of LULL1 compared to LAP1 in mouse liver. Although immunoblotting does not allow direct comparison of the expression level of two proteins, we detected robust expression of LULL1 and LAP1 in wild type mouse liver tissue and hepatocyte lysates. Further examination of mRNA expression showed similar levels of those encoding LAP1B and LAP1C, two LAP1 isoforms, and LULL1 in livers from wild type mice at 4 months of age (Supplementary Table 2). Mouse ENCODE transcriptome data also showed similar expression of the two genes in adult liver (LAP1B: RPKM 5.7, LULL1: RPKM 6.5) ^16^. These data indicate that the minimal effects of LULL1 depletion on lipid secretion and accumulation are not likely explained by large differences in protein expression levels. The absence of excess lipid accumulation in hepatocytes lacking LULL1 suggests that LAP1 may sufficiently stimulate torsinA activity in the absence of LULL1. However, the more-modest steatosis observed in mice with hepatocyte-specific depletion of LAP1 compared to those with depletion of torsinA suggests that LULL1 alone nonetheless provides partial torsinA function when present in cells.

### Depletion of both LAP1 and LULL1 from mouse hepatocytes leads to steatosis similar to that occurring with torsinA depletion

We hypothesized that if LAP1 and LULL1 are the only or major activators of torsinA in hepatocytes, depletion of both will phenocopy the marked hepatic steatosis and VLDL secretion defect in A-CKO mice with depletion of torsinA from hepatocytes. To generate combined depletion of both proteins, we used shRNA-mediated gene depletion, as we could not delete the genes encoding both LAP1 and LULL1 by crossing mouse lines due to their tight physical linkage in the genome ^4^. We designed four shRNAs to deplete LAP1 and selected one with the highest knockdown efficiency in a mouse cell line (Supplementary Fig. 3a). We then generated an adenoviral vector to express the shRNA under the control of a U6 promoter (Ad-Lap1) for *in vivo* depletion of LAP1 from mouse hepatocytes. To test the efficiency of Ad-Lap1shRNA, we transduced cultured wild type primary hepatocytes with this or a similar construct expressing a control scrambled shRNA (Ad-ctrl) and confirmed the depletion of LAP1 48 hours later by immunoblotting (Supplementary Fig. 3b and 3c). Accordingly, immunofluorescence microscope studies showed no nuclear rim staining of LAP1 in the hepatocytes transduced with Ad-Lap1shRNA (Supplementary Fig. 3d). We then intravenously administrated the adenoviral constructs to wild type mice to test *in vivo* efficacy of LAP1 depletion. Analysis of liver sections by immunofluorescence microscopy 5 days post injection demonstrated depletion of LAP1 (Supplementary Fig. 3e).

After confirming the efficacy of Ad-Lap1 for depleting LAP1, we administered it (or Ad-ctrl) intravenously to LUL-flox and LUL-CKO mice. Immunoblotting of whole liver lysates showed that LUL-flox and LUL-CKO mice treated with Ad-Lap1 had reductions in expression in all LAP1 isoforms, with a somewhat greater decrease in LAP1C in the LUL-flox mice (Fig. 3a). Quantification of total LAP1 signals on immunoblots of whole liver lysates showed that Ad-Lap1 treatment produced approximately 50% reductions in LAP1 expression in LUL-flox and LUL-CKO mice, and a slight but non-significant trend towards increased torsinA expression in LUL-CKO mice (Fig. 3b). On gross inspection 5 days after adenoviral vector treatment, livers from LUL-flox mice given Ad-ctrl or Ad-LAP1 and from LUL-CKO mice administered Ad-ctrl were grossly normal, whereas livers from LUL-CKO mice treated with Ad-Lap1, which have depletion of both LAP1 and LULL1 from hepatocytes, were grossly white, similar to livers from A-CKO mice with hepatocyte-specific depletion of torsinA (Fig. 3c). Histological examination of sections stained with hematoxylin and eosin showed increased fat content in livers from mice with depletion of both LAP1 and LULL1 from hepatocytes – similar to that in livers from A-CKO mice with depletion of torsinA – and a more modest increase in fat in mice with depletion of only LAP1 from hepatocytes (Fig. 3d). A pathologist blind to genotype and treatment invariably gave a steatosis score of 3 (on a scale of 0 to 3) to every section examined from livers of mice with combined depletion of LAP1 and LULL1 or with depletion of torsinA alone; the steatosis was mostly microvesicular. Micrographs of liver sections stained with Oil Red O confirmed significant fat in livers of mice with depletion of both LAP1 and LULL1 from hepatocytes, which was similar to those from A-CKO mice with depletion of torsinA (Fig. 3e). Livers from mice with hepatocyte-specific depletion of both LAP1 and LULL1 and from A-CKO mice with depletion of torsinA had significantly increased TG content compared to controls and mice with depletion of only LAP1 from hepatocytes (Fig. 3f). These results show that depletion of both known torsinA activators – LAP1 and LULL1 – from hepatocytes produce a dramatic degree of liver steatosis similar to what occurs with depletion of torsinA.

**Fig. 3.**
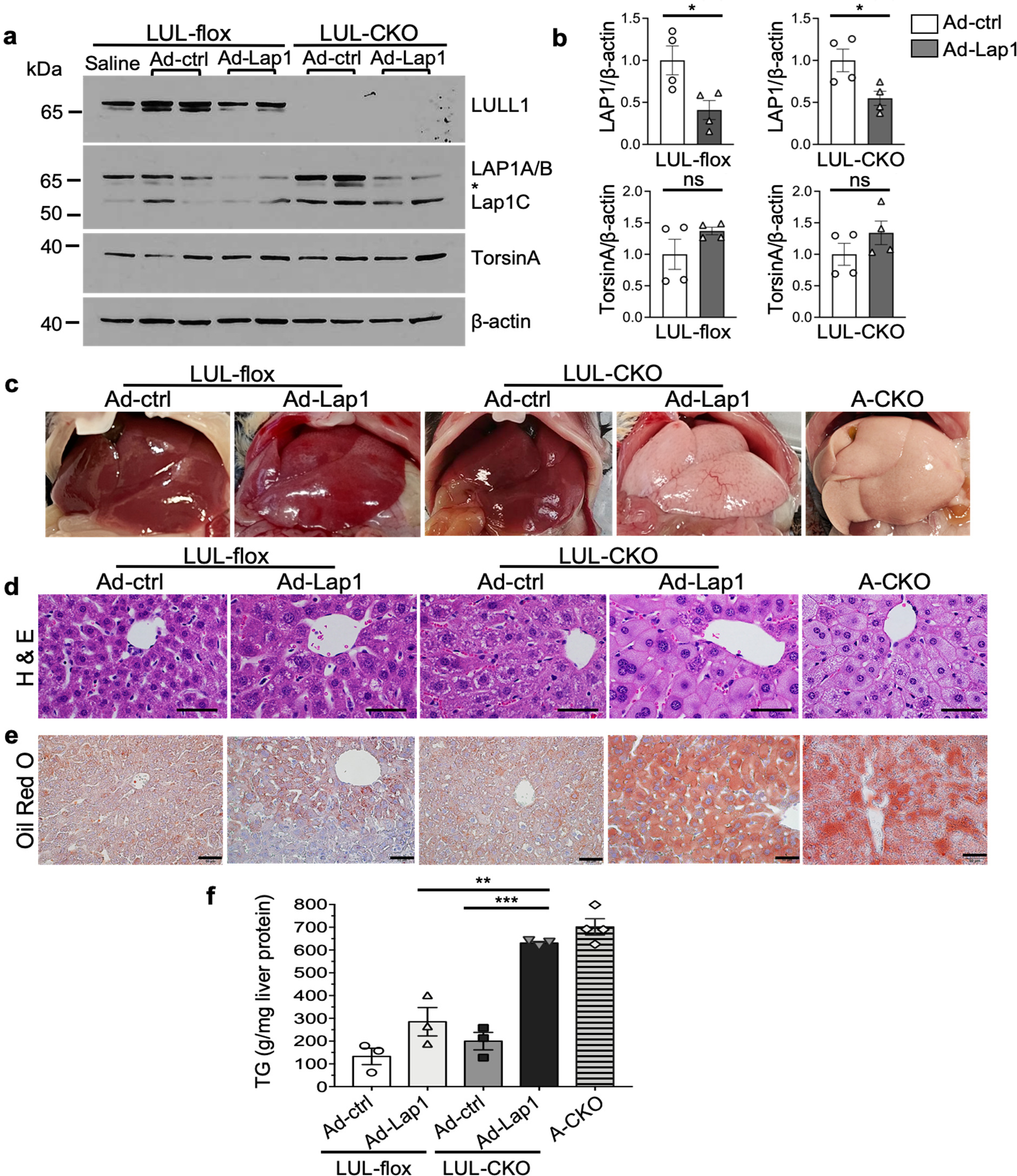
Depletion of LAP1 and LULL1 from hepatocytes leads to liver steatosis similar to that occurring with torsinA depletion. **a** Immunoblots of liver lysates from LUL-flox and LUL-CKO mice administered adenoviral vectors expressing either control scrambled shRNA (Ad-ctrl) or shRNA that targets LAP1 (Ad-Lap1). Lysates from liver of a LULL-flox mouse treated with saline is also included as a control. Antibodies against LULL1, LAP1, torsinA and β-actin were used. *Indicates non-specific bands often detected in whole liver lysates using polyclonal antibodies against LAP1. **b** Relative ratios of band densities of LAP1 (all isoforms) or torsinA to β-actin in blots as shown in panel. Band densities were measured from two independent immunoblots using protein extracts from two mice per group each. Values are means ± SEM with each circle or triangle representing the value from each immunoblot (n = 4). \**p* < 0.05, ns = not significant by Student’s t-test. **c** Representative photographs of livers from LUL-flox and LUL-CKO mice injected with Ad-ctrl or Ad-Lap1. Photograph of a liver from an A-CKO mouse with depletion of torsinA from hepatocytes is shown for comparison. **d** Representative light photomicrographs of liver sections stained with hematoxylin and eosin (H & E) from LUL-flox and LUL-CKO mice administered with Ad-Lap1 or Ad-ctrl. Section from an A-CKO mouse is shown for comparison. Scale bars: 50 μm. **e** Representative light photomicrographs of liver sections stained with Oil Red O from LUL-flox and LUL-CKO mice injected with adenoviruses expressing either Ad-ctrl or Ad-Lap1. A Section from an A-CKO mouse is shown for comparison. Scale bars: 50 μm. **f** TG content in livers from LUL-flox and LUL-CKO mice administered Ad-ctrl or Ad-Lap1. Mice were fasted for 4-5 hours before isolating livers. Liver TG values from 4 A-CKO mice were included for comparison. Values are means ± SEM with each circle or triangle representing the value from an individual mouse (n = 3-4). \*\**p* < 0.01, \*\*\**p* < 0.001 by ANOVA.

Whole liver lysates contain proteins from cells other than hepatocytes. We therefore isolated hepatocytes from livers of LUL-flox and LUL-CKO mice 5 days after injection of the adenoviral vectors to better assess the depletion of LAP1. Immunoblotting of isolated hepatocytes demonstrated diminished expression of all LAP1 isoforms in hepatocytes isolated from both LUL-flox and LUL-CKO mice administered shRNA Ad-Lap1; however, there was an apparently greater decrease in LAP1A/B isoforms in LUL-CKO mice (Fig. 4a). Quantification of total LAP1 signals on immunoblots of hepatocyte lysates showed that Ad-Lap1 treatment produced approximately 75% to 85% reductions in LAP1 expression in LUL-flox and LUL-CKO mice and a significant increase in torsinA expression (Fig. 4b). The measured decreases in LAP1 expression and increases in torsinA expression in isolated hepatocytes were of greater magnitude than when measured in whole liver lysates. While residual expression of LAP1 was detected by immunoblotting, we observed little to no nuclear rim labeling with anti-LAP1 antibodies in confocal immunofluorescence micrographs of hepatocytes from both LUL-flox and LUL-CKO mice that received Ad-Lap1 (Fig. 4c). We previously showed that torsinA depletion from hepatocytes leads to the accumulation of mostly microvesicular lipid droplets in the cytosol, whereas LAP1 depletion leads to nuclear lipid droplets with more modest cytosolic lipid accumulation ^11, 15^. We therefore stained hepatocytes with BODIPY and DAPI to analyze the subcellular localization of lipid droplets in those lacking LAP1 and LULL1 (Fig. 4d). While we observed relatively low levels of normal shaped lipid droplets in hepatocytes from LUL-flox and LUL-CKO mice treated with Ad-ctrl, there was increased cytosolic lipid and nuclear lipid droplets in hepatocytes from LUL-flox mice treated with Ad-Lap1 to deplete LAP1. In hepatocytes from LUL-CKO mice treated with Ad-Lap1, resulting in depletion of both LULL1 and LAP1, there was striking accumulation of cytosolic lipids with minimal to no nuclear lipid droplets similar to that in hepatocytes from A-CKO mice with depletion of torsinA from hepatocytes (Fig. 4d).

**Fig. 4.**
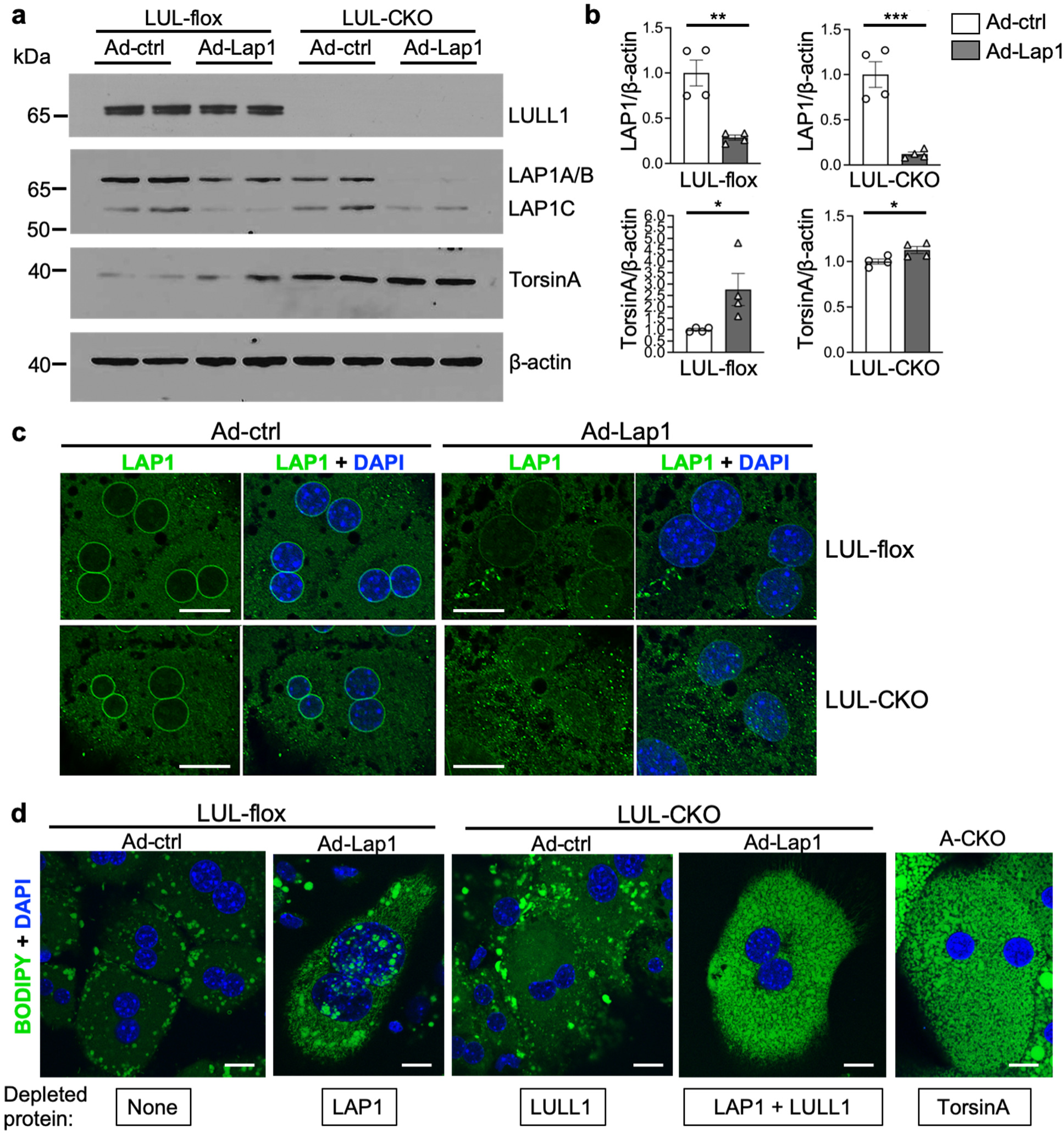
Depletion of LAP1 and LULL1 from hepatocytes leads to lipid droplet distribution similar to that occurring with torsinA depletion. **a** Immunoblots of hepatocyte lysates from LUL-flox and LUL-CKO mice administered adenoviral vectors expressing either control scrambled shRNA (Ad-ctrl) or shRNA that targets LAP1 (Ad-Lap1). Antibodies against LULL1, LAP1, torsinA, and β-actin were used. **b** Relative ratios of band densities of LAP1 (all isoforms) or torsinA to β-actin blots shown in panel a. Band densities were measured from two independent immunoblots using protein extracts from two mice per group each. Values are means ± SEM with each circle or triangle representing the value from each immunoblot (n = 4). \**p* < 0.05, \*\**p* < 0.01, \*\*\**p* < 0.001 by Student’s t-test. **c** Representative immunofluorescence confocal photomicrographs of hepatocytes from LUL-flox and LUL-CKO mice administered Ad-ctrl or Ad-Lap1. Cells were stained with anti-LAP1 antibody and DAPI (blue). Scale bars: 20 μm. **d** Representative immunofluorescence confocal micrographs of hepatocytes from LUL-flox and LUL-CKO administered Ad-ctrl or Ad-Lap1. Cells were stained with BODIPY (green) and DAPI (blue). A torsinA deficient hepatocyte from an A-CKO mouse is shown for comparison. Scale bars: 10 μm. Proteins depleted from the cells are indicated at the bottom of each image.

We performed additional confocal microscopic analyses to quantify nuclear lipid droplets in hepatocytes lacking LAP1, LULL1, or both (Fig. 5a). While almost all hepatocytes with depletion of only LAP1 had more than one nuclear lipid droplet with approximately 30% having 10 or more per nucleus, few hepatocytes with depletion of both LAP1 and LULL1 had more than 10 lipid droplet per nucleus (Fig. 5b). In hepatocytes with depletion of only LAP1, we found that 61± 7% (mean ± SEM) of nuclei contained had 5 or more nuclear lipid droplets, whereas only 13.2 ± 2% hepatocytes with depletion of both LAP1 and LULL1 had 5 or more nuclear lipid droplets. (Fig. 5c). These results indicate that depletion of LAP1 and LULL1 from hepatocytes produced the same subcellular distribution of lipids that occurs in hepatocytes with loss of torsinA.

**Fig. 5.**
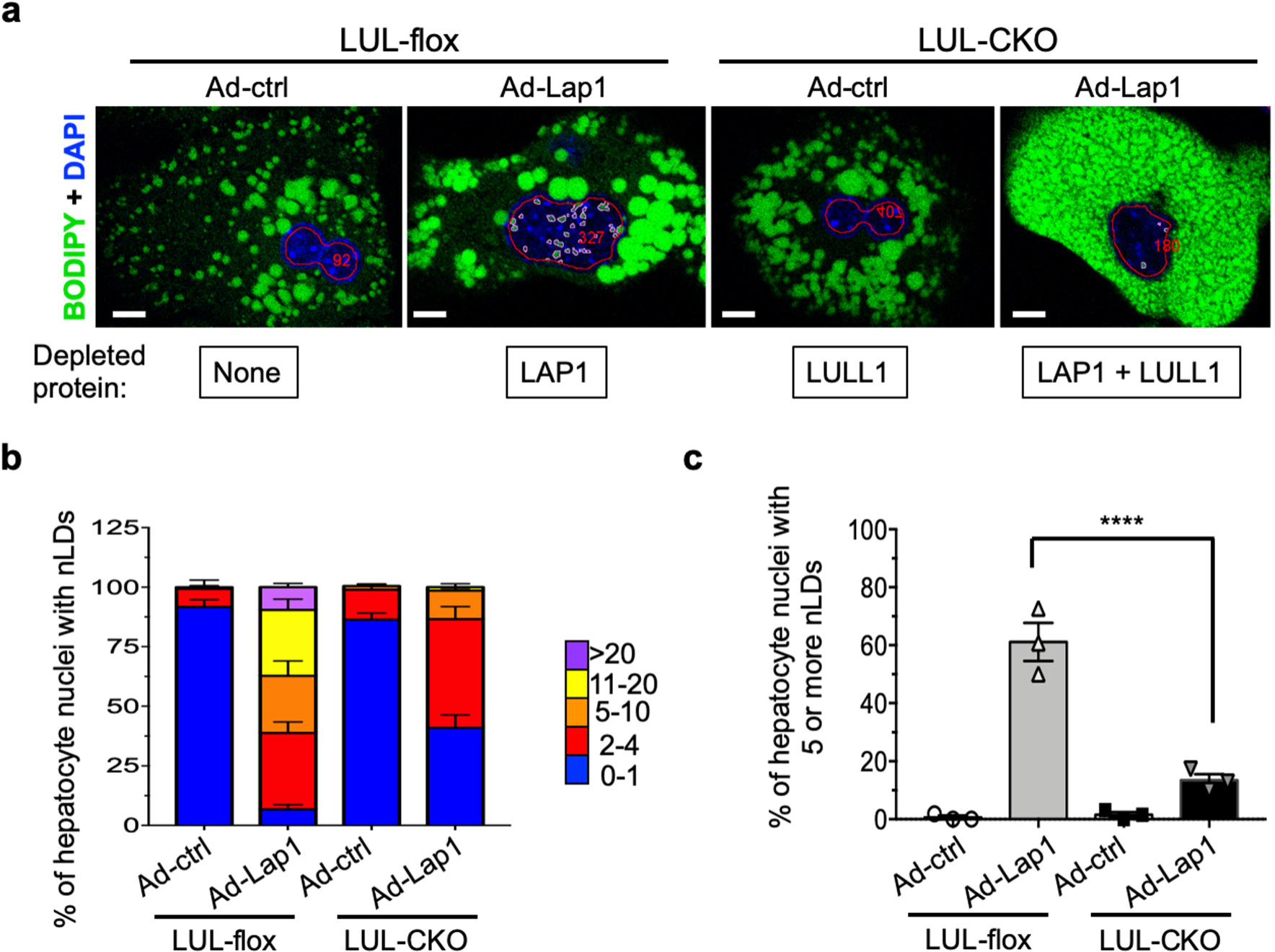
Nuclear lipid droplets in hepatocytes with depletion of LAP1, LULL1 or both. **a** Representative immunofluorescence confocal micrograph of BODIPY (green) and DAPI (blue) stained hepatocyte cultures of LUL-flox and LUL-CKO mice injected with the adenovirus expressing either Ad-ctrl or Ad-Lap1. Red lines mark nuclei and pink lines nuclear lipid droplets. Red numbers indicate the cell count according to the automated scoring method (see the method section). Scale bars: 10 μm. Proteins depleted from the cells are indicated at the bottom of each image. **b** Stacked column graphs with different colors representing the percentages of hepatocyte nuclei containing the indicated numbers of nuclear lipid droplets (nLDs). We analyzed a total of 131 (LUL-flox mouse administered Ad-ctrl), 101 (LUL-flox mouse administered Ad-Lap1), 136 (LUL-CKO administered Ad-ctrl), and 94 (LUL-CKO mouse administered Ad-Lap1) nuclei of hepatocytes cultured on three different coverslips (n=3 per group). The values within the multicolored bars are means + SEM. **c** Percentage of hepatocyte nuclei containing more than 5 nLDs calculated from data shown in b. Values are means ± SEM with each circle or triangle the value from each coverslip (n = 3). \*\*\*\**p* < 0.0001 by ANOVA.

### Depletion of both LAP1 and LULL1 from mouse hepatocytes leads to reduced liver VLDL secretion similar to that occurring with torsinA depletion

We next tested if steatosis in livers of mice with hepatocytes deficient in both LAP1 and LULL1 is caused by defective VLDL secretion, as is the case in A-CKO mice with depletion of torsinA from hepatocytes ^11^. Due to the transient effect of adenoviral-mediated gene delivery systems, we performed *in vivo* TG secretion assays in LUL-flox or LUL-CKO mice 5 days after administration of either Ad-ctrl or Ad-Lap1 (Fig. 6a). We observed no significant changes in TG concentration between groups with no protein depleted (LULL-flox injected with Ad-ctrl) or only LULL1 depleted (LULL-CKO injected with Ad-ctrl) at any timepoint. In contrast, we observed significant reductions of TG secretion rate in mice with LAP1 single depletion (LUL-flox mice injected with Ad-Lap1) and further reduction in the group with combined LAP1 and LULL1 depletion (LUL-CKO mice injected with Ad-Lap1) as compared to groups with no protein depleted or only LULL1 depleted (Fig. 6b). Plasma proteins collected at 60 minutes after injection of ^35^S-methionine with tyloxapol were analyzed by SDS-PAGE followed by autoradiography to detect radiolabeled apolipoprotein apo B100 or apoB48 (Fig. 6c) Analysis of newly synthesized proteins entering the plasma after injection of ^35^S-methionine showed a significant decrease of apoB100 in the LAP1-depleted group (LUL-flox mice injected with Ad-Lap1) and further decrease of apoB100 in the group with combined depletion of LAP1 and LULL1 (LUL-CKO mice injected with Ad-Lap1) compared to mice with no protein depleted; the same trend did not meet statistical significance for apoB48 (Fig. 6d). These results indicate that hepatocytes with combined depletion of LAP1 and LULL1 accumulate large amounts of lipids in association with VLDL secretion defects, which is similar to what we observed in A-CKO mice with depletion of torsinA from hepatocytes ^11^.

**Fig. 6.**
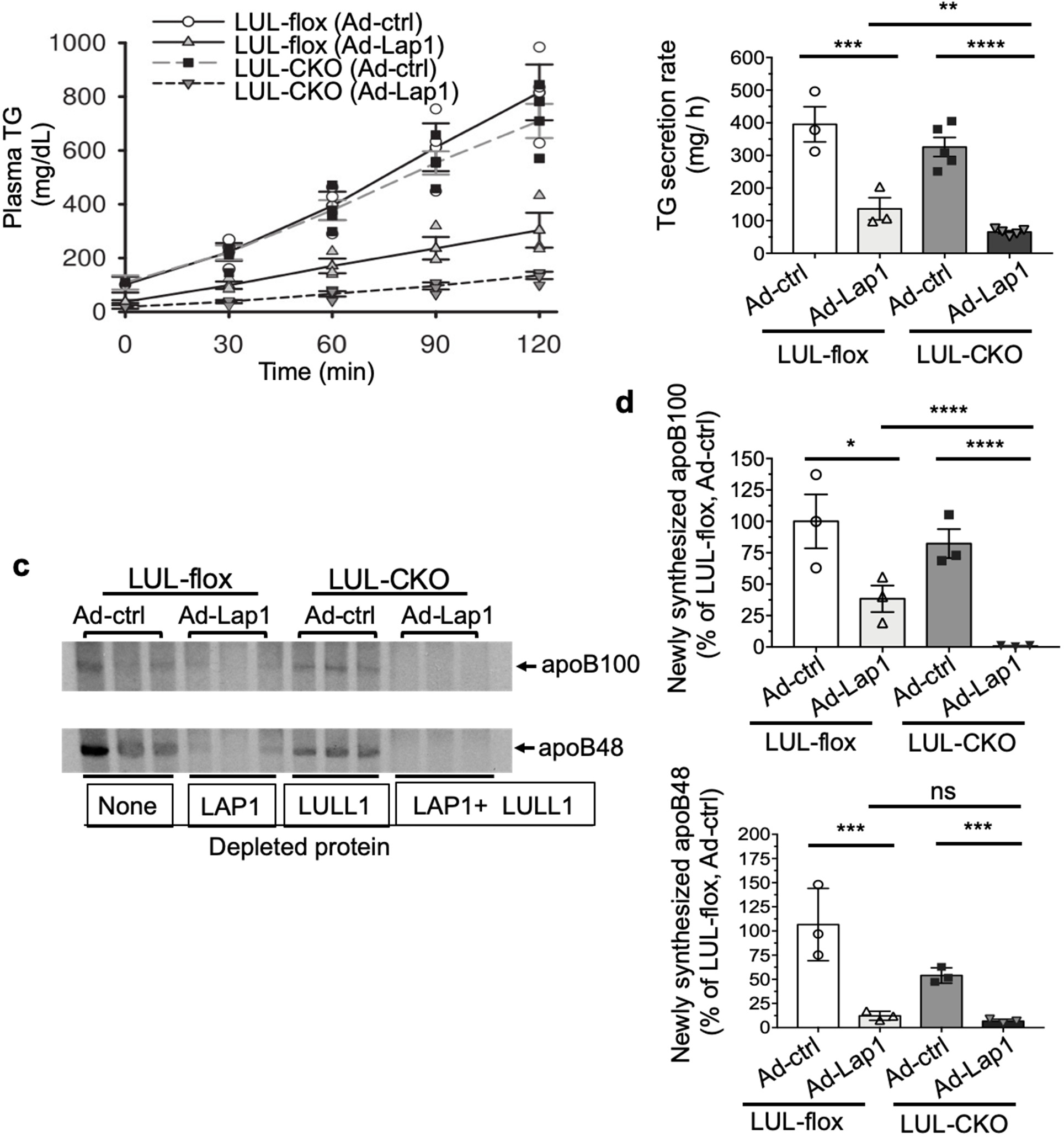
Depletion of LAP1 and LULL1 from hepatocytes leads to decreased liver TG secretion similar to that occurring with torsinA depletion. **a** Plasma TG concentrations versus time after injection of tyloxapol to block peripheral uptake in LUL-flox and LUL-CKO mice 5 days post injection of adenoviral vectors expressing either control scrambled shRNA (Ad-ctrl) or shRNA that targets LAP1 (Ad-Lap1). Values are means ± SEM with each circle, square or triangle representing the value for an individual mouse (n = 3-5). **b** TG secretion rates were calculated by changes in plasma concentrations between the 30 and 120 minute time points in a. **c** Autoradiogram of SDS-polyacrylamide gel showing ^35^S-labeled plasma proteins collected 60 minutes after injection of mice with ^35^S-methionine. Each lane shows protein from an individual mouse (n=3 per group). Migration of ^35^S-labeled apoB100 and apoB48 is indicated. Proteins depleted from the cells are indicated at the bottom of the autoradiogram. **d** Bands corresponding to apoB100 (upper panel) and apoB48 (lower panel) as shown in c were quantified by densitometry and shown as a % of the value from the Ad-ctrl injected LUL-flox group. In panels b and d, values are means ± SEM with each circle, square or triangle representing the value from an individual mouse (n = 3-5). \**p* < 0.05, \*\**p* < 0.01, \*\*\**p* <0.001, \*\*\*\**p* <0.0001, (ns) = not significant by ANOVA after raw values were log-transformed as values from a TG secretion timepoint showed heteroscedasticity by Levene’s test and violation of normality by Shapiro-Wilk test (see Statistics section).

## Discussion

Several previous studies have demonstrated direct interactions between torsinA and LAP1 and LULL1 using various cell biological, biochemical and structural approaches ^4–6, 9^. However, neither the physiological relevance of these interactions nor the relative contribution of the two activators to torsinA function *in vivo* has been demonstrated. We now show that combined depletion of LAP1 and LULL1 from hepatocytes leads to marked hepatic steatosis and VLDL secretion defects similar in magnitude to that occurring in A-CKO mice with depletion of torsinA from hepatocytes. These results provide compelling evidence that LAP1 and LULL1 both modulate torsinA function in controlling hepatocyte fat content *in vivo*.

LAP1 depletion from hepatocytes causes decreased VLDL section and steatosis; however, these are less severe than when torsinA is depleted ^11^. This led us to hypothesize that LULL1 can partially compensate for LAP1 in liver lipid metabolism. However, LULL1 depletion from hepatocytes does not cause hepatic steatosis or decreased VLDL secretion. The interaction of LAP1 and LULL1 with torsinA to restore its function in hepatocytes is therefore more complex than a binding to two different proteins, expressed at similar levels, with similar avidities. One possibility is that there is an unrecognized interaction with LAP1 that is essential for LULL1 to fully establish torsinA function. As LAP1 is a transmembrane protein of the inner membrane of the nuclear envelope, another possibility is that torsinA function in the perinuclear space is important for hepatocyte VLDL secretion. LULL1 is a monotopic transmembrane protein with a cytoplasmic domain of 215 amino acids ^4^. It is localized throughout the entire ER, which includes the outer nuclear membrane. Given the size of its cytoplasmic domain, some fraction LULL1 can diffuse through the interconnected ER, outer nuclear membrane, pore membrane and inner nuclear membranes ^17^. Therefore, in the absence of LAP1, some LULL1 is localized to the nuclear envelope and may be responsible for providing the partial function of torsinA there. If so, it raises the question of why nuclear envelope-localized LAP1 appears to be the major activator of torsinA in hepatic lipid secretion despite LULL1 being localized to VLDL-forming components in the more peripheral ER.

The luminal regions of LAP1 and LULL1 contain a conserved domain where the arginine finger of canonical AAA+ ATPases is found ^18^. The interaction of one of these activators with torsinA is predicted to generate an active site for ATPase activity. LAP1 or LULL1 binding to torsinA *in vitro* stimulates ATPase activity ^5, 6^. However, we cannot apply the assays used *in vitro* to liver or hepatocyte lysates because a substrate for torsinA enzymatic activity has not been identified. It therefore remains unknown if ATPase activity of torsinA and its activators or some other process underlies their function in VLDL secretion and liver lipid metabolism.

A significant number of nuclei in hepatocytes with depletion of LAP1 contain nuclear lipid droplets, which are rarely observed with depletion of torsinA. Nuclear lipid droplets in hepatocytes with depletion of LAP1 likely arise from invaginations of the inner nuclear membrane, which are also known as type 1 nucleoplasmic reticulae ^15^. Nuclear lipids associated with type I nucleoplasmic reticula also occur in hepatocellular carcinoma-derived cell lines and they are increased by depletion of SUNs, which like LAP1 are integral proteins of the inner nuclear membrane ^19, 20^. However, hepatocytes with combined depletion of LAP1 and LULL1 more closely resemble hepatocytes with depletion of torsinA, with few if any nuclear lipid droplets but numerous small and some larger cytoplasmic ones. Electron microscopy has shown that the numerous small lipid droplets in hepatocytes with depletion of torsinA appear to be arranged in tubular configurations, suggesting that they may be in the ER ^11^. This suggests that depletion of both LAP1 and torsinA, which presumably leads to loss of most torsinA function in hepatocytes, causes a defect in the ER that somehow “overrides” the generation of nuclear lipid droplets.

Our studies demonstrate that torsinA and its activators LAP1 and LULL1 profoundly influence VLDL secretion and hepatic steatosis. Deciphering the downstream mechanisms connecting loss of function of torsinA and its activators to the pronounced hepatic phenotypes will be the focus of future research. It is also unclear how these proteins may related to human lipid metabolism or conditions such as steatotic liver disease. One study has tentatively linked serum TG concentrations to polymorphisms in *TOR1AIP1* encoding LAP1 ^21^. An intronic variant in *TOR1B* encoding torsinB, a paralogue of torsinA, has also been linked to steatotic liver disease ^22^. Given the profound phenotypes we have observed in mice, polymorphisms or mutations in the gene encoding torsinA, LAP1, and LULL1 may influence hepatic and lipid metabolic traits in humans.

## Methods

### Mice

The Columbia University Institutional Animal Care and Use Committee approved the protocols and procedures. We generated mice with hepatocyte-specific depletion of LULL1 by crossing female mice with homozygous floxed alleles of *Tor1aip2* encoding LULL1 to male Alb-Cre transgenic mice (Jackson Laboratory stock number 003574) (see Supplementary Figure 1). Genotyping was performed using the method described in Supplementary Figure 1 and Supplementary Table 1. L-CKO mice with depletion of LAP1 from hepatocytes, A-CKO mice with depletion of torsinA from hepatocytes and mice with floxed alleles of *Tor1a* encoding torsinA have been previously described ^11, 23^. The genetic background of all mice was C57BL/6J. Mice were housed in a climate-controlled room with a 12-hour light/12-hour dark cycle and fed regular chow diet (Purina Mills 5053). Both male and female mice were used for experiments.

### Primary hepatocyte isolation

Primary hepatocytes were isolated according to previously described methods ^24^. Briefly, mice were perfused with Hanks balanced salt solution without calcium (Thermo Fisher Scientific) with 8 mM HEPES (Thermo Fisher Scientific) via the abdominal inferior vena cava after cutting the portal vein to allow outflow of the perfusate. They were perfused for 8 minutes at a rate of 5 ml/minute at 37°C. This was followed by perfusion at the same rate of DMEM (Thermo Fisher Scientific) with 80 mg/100 ml of collagenase type I (Worthington Biochemical) for 6 minutes. The liver was removed and minced in a Petri dish containing 4 ml of the same warm DMEM collagenase mixture for an additional 2 to 4 minutes. Ice-cold DMEM was added and the digested tissue was filtered through a nylon mesh and collected in a 50 ml conical tube. The suspension was centrifuged for 5 minutes at 500 rpm. The supernatant was aspirated and the cell pellet was washed 3 times with 30 ml of ice-cold DMEM. Viable cells were counted after staining with trypan blue and were >90%. Isolated primary hepatocytes were used for experiments within 24 or 48 hours after isolation. In some cases, isolated hepatocytes from wild type mice were frozen using Cryo-JIN solution (Revive Organtech), a specialized freezing media for primary hepatocytes, and thawed for immunofluorescence staining according to the manufacturer’s instructions.

### Cell culture

C2C12 cells were purchased from ATCC (CRL-1772). Culture conditions were according to ATCC protocols. The isolated primary mouse hepatocytes were plated onto collagen-coated 6-well plates at a density of 500,000 cells/well in 4 ml of DMEM (Thermo Fisher Scientific) plus 10% fetal bovine serum, 1% penicillin-streptomycin (Thermo Fisher Scientific), and 1% HEPES, and cultured for 18 to 48 hours for subsequent experiments including immunoblotting and apoB secretion assays.

### Generation of adenoviral vectors and administration to mice

To generate an adenoviral vector expressing shRNA targeting LAP1, four different shRNA sequences targeting mouse LAP1 were synthesized and cloned into a pSuper-retro-puro plasmid (Oligoengine) according to the manufacturer’s instructions, Knockdown efficiency was tested by transfected C2C12 cells and immunoblotting lysates. After choosing the sequence showing most efficient knockdown (CAAGTGTCTGAGTGAACAAAT corresponding to a portion of *Tor1aip1* exon 10), a synthesized oligonucleotide was inserted under the U6 promoter in an adenoviral vector, and virus was packaged, amplified, and purified (Welgene). A control adenoviral vector expressing a scrambled shRNA under control of the same promoter was purchased from Welgene. Adenovirus was delivered by retro-orbital injections in 3-6 months old mice at doses of 1-2.5 x 10^9^ plaque-forming-units/kg.

### Immunoblotting

Livers and cultured cells were homogenized in cell lysis buffer (Cell Signaling) supplemented with 0.1% SDS, 0.2% sodium deoxycholic acid, Protease Inhibitor Cocktail (Sigma-Aldrich), and 1 mM phenylmethylsulfonyl fluoride (Sigma-Aldrich). Proteins in the samples were denatured by boiling in Laemmli sample buffer ^25^ containing β-mercaptoethanol for 5 minutes, then separated by 10% or 4% SDS-PAGE, and transferred to nitrocellulose membranes. For immunoblotting, membranes were washed with blocking buffer (tris buffer saline (TBS) containing 5% nonfat dry milk and 0.1% polysorbate-20) for 30 minutes and then probed with primary antibodies in blocking buffer overnight at 4°C. Primary antibodies were against LAP1 ^4^, LULL1 ^4^, torsinA (Abcam, ab34540), β-actin (Santa Cruz Biotechnology, sc-47778), myosin heavy chain (Santa Cruz Biotechnology, C-20641), γ-tubulin (Sigma-Aldrich, T5326) and GAPDH (Ambion, AM4300). Membranes were washed with TBS containing 0.1% polysorbate-20 and then incubated in blocking buffer containing horseradish peroxidase-conjugated secondary antibodies (GE Healthcare) for 1 hour at room temperature. After washing with TBS containing 0.1% polysorbate-20, membranes were soaked with Enhanced Chemiluminescence substrates (Thermo Fisher Scientific) and exposed to X-ray films. To quantify signals, the exposed films were scanned and the densities of the bands were quantified using ImageJ software ^26^.

### Histopathology

Livers were immediately excised and blotted dry after euthanizing mice. Portions of excised livers were fixed in 10% formalin for 48 hours. Paraffin block preparation, sectioning into 5 µm slices and staining with hematoxylin and eosin were performed by the Histology Service of the Columbia University Molecular Pathology Core. Other portions of livers were transferred to a 30% sucrose solution and frozen sections 5 µm think sections were prepared by the Histology Service of the Columbia University Molecular Pathology Core for staining with Oil Red-O (Polyscience #25962). Stained sections were photographed using a BX53 upright microscope attached to a DP72 digital camera (Olympus). A liver pathologist (M.J.L.) blind to genotype or treatment analyzed the hematoxylin and eosin-stained sections and provided a steatosis score from 0 (<5% of parenchymal involvement) to 3 (>66%) considering total macrovesicular and microvesicular fat.

### Quantification of liver TG

Liver lipids were extracted with a modified Folch method as previously described ^27^. Briefly, a snap-frozen piece of liver (∼100 mg) was homogenized in PBS and lipids were first extracted with chloroform:methanol (2:1 v/v) and a second time with chloroform:methanol:water (86:14:1 v/v/v). The organic layer was dried under nitrogen gas and resolubilized in chloroform with 15% Triton X-100 (Sigma-Aldrich). It was dried under nitrogen gas and finally brought up in double-distilled H_2_O.

### *In vivo* TG and apoB secretion

*In vivo TG* and apoB secretion rates were determined as previously described ^24^. Briefly, mice were fasted for 4 to 5 hours prior to an intravenous injection of a mixture of 200 μCi ^35^S-methionine and 500 mg/kg tyloxapol (Sigma-Aldrich, T8761-50G) in 0.9% NaCl. Tyloxapol inhibits both the lipolysis and tissue uptake of lipoproteins in mice, and the subsequent concentrations of TGs and ^35^S-apoB in plasma can be used to calculate their rates of secretion. Blood samples were collected before injection and 30, 60, 90, and 120 minutes after injection of tyloxapol and ^35^S-methionine. Plasma TG concentrations were measured using a colorimetric assay (Wako Diagnostics, 461-08992). For apoB secretion, whole plasma samples from the 60 minute time point were subjected to 4% SDS-PAGE. The volumes of the samples were adjusted as determined by trichloroacetic acid precipitable ^35^S-labeled plasma proteins. The gel was dried and exposed to x-ray film to quantitate labeled apoB proteins by densitometry.

### Fluorescence microscopy

For immunofluorescence microscopy of whole livers, frozen sections were post-fixed in 4% paraformaldehyde for 5 minutes at room temperature, then washed with PBS 3 times. Sections were then subjected to treatment with Antigen Unmasking Solution (Vector Laboratory, #H-3300) according to the manufacturer’s instructions. The treated sections were incubated with blocking solution (5% bovine serum albumin, 2% normal goat serum, and 0.5% Triton X-100 in PBS) for 30 minutes at room temperature and subsequently incubated with anti-LAP1 antibody ^4^ in the blocking solution for overnight at 4°C. After 3 washing with PBS containing 0.1% polysorbate-20, the sections were incubated with Alexa fluor 488 conjugated secondary antibody (Thermo Fisher Scientific) with DAPI for 1 hour at room temperature. After 3 additional washes, sections were mounted with ProLong mounting media (Thermo Fisher Scientific). Widefield fluorescent images were acquired using a BX53 upright fluorescent microscope attached to a DP72 microscopy.

For fluorescence microscopy of isolated hepatocytes, cells were seeded onto the collagen-coated coverslips in 6-well plates (150,000 cells/well) and grown over night in a 37°C and 5% CO_2_ incubator before fixation or lipid staining. Cultured primary hepatocytes on collagen-coated cover slips were stained with 2 mM BODIPY (Thermo Fisher Scientific, D-3922) in Hank’s Balanced Salt Solution (Thermo Fisher Scientific) for 30 minutes. Subsequently, cells were fixed with 4% paraformaldehyde for 15 minutes at room temperature, then washed and stained with DAPI. For antibody staining, the fixed cells were incubated with primary antibodies overnight. Primary antibodies used were anti-LAP1 ^4^, anti-lamin A/C (Abcam, ab133256) and anti-torsinA (Abcam, ab34540). After incubation with primary antibodies, cells were washed and then labeled with Alexa488-conjugated goat anti-mouse and rhodamine-conjugated goat anti-rabbit secondary antibodies (Jackson ImmunoResearch Laboratories). Confocal micrographs were obtained using a TCS SP8 DLS microscopy (Leica) in the Medicine Microscopy Core at Columbia University Irving Medical Center. For images shown in Fig. 2d, we used a Nikon A1 RMP microscope using NIS-elements software (Nikon) at the Confocal and Specialized Microscopy Shared Resource of the Herbert Irving Comprehensive Cancer Center at Columbia University Irving Medical Center. Acquired fluorescent images were processed using NIS-elements software, Image J or Fiji ^26^.

For automated analysis of fluorescence images, each batch of hepatocytes was plated on 3 coverslips and lipid droplets stained with BOPIDY 493/502 (Thermo Fisher Scientific) according to the manufacturer’s instructions. DAPI was included in the media for nuclear staining. Fluorescence images from 5-10 different regions were captured. Cytosolic and nuclear lipid droplets were quantified using macro programs created in Fiji/Image J ^26^. For confocal images, DAPI stained nuclei were subject to Gaussian blur and triangle thresholding, and resulting nuclei objects were reduced in size by 1.8 microns to exclude cytosolic florescence. BODIPY stained lipid droplets within nuclear objects were subject to a spot enhancing filter ^28^, Gaussian blur filter, and triangle threshold to create droplet objects. Nuclei and lipid droplet object outlines were overlaid on to fluorescence images, cells were counted, parameters were measured, and a data table was created containing nuclear area, nuclear region, nuclear lipid droplet area, and nuclear lipid droplet BODIPY fluorescence intensity. For cytosolic fluorescence measurements, the analysis macro was modified to include object creation of individual cells using the deep learning software, CellPose ^29^, which was run within ImageJ. Nuclei objects found to reside within the cytosol were subtracted from the cytosol objects using the XOR function within ImageJ ROI manager and cytosol area and BOPIDY intensity were output to a table.

### Statistics

Statistical methods used are described in the legend of each relevant figure. Unpaired 2-tailed Student’s t-test was used to compare differences between any two groups (GraphPad Prism, version 9.4.0). Comparisons for four groups were performed by using one-way ANOVA with a subsequent Tukey post-hoc test. Before proceeding with ANOVA, the fundamental assumptions of normality and homoscedasticity were assessed to ensure the validity of the statistical tests. The normality assumption was assessed using the Shapiro-Wilk test. A *p*-value < 0.05 indicated a violation of normality. In such cases, data transformation was performed to address non-normality. The assumption of homoscedasticity was evaluated using Levene’s test. A *p*-value < 0.05 indicated the presence of heteroscedasticity, which required corrective measures to achieve homogeneity of variances. If either or both the Shapiro-Wilk test and Levene’s test reported a *p*-value < 0.05, a log transformation was applied to the data to rectify violations of the ANOVA assumptions. After log transformation, the Shapiro-Wilk test and Levene’s test were re-administered to verify the successful normalization of the data and homogenization of variances. Once the assumptions of normality and homoscedasticity were met following the data transformation, the one-way ANOVA with Tukey post-hoc test was applied to identify specific pairwise differences between the group means. To validate the effectiveness of the log transformation, the ANOVA residuals plot was examined both before and after the transformation, allowing for visual confirmation of the presence of non-normality and heteroscedasticity before the transformation and their absence after. Statistical analyses for multiple group comparisons were performed using Statistical Analysis Software, version 9.4M6 (SAS Institute Inc). A significance level of α = 0.05 was adopted to determine the statistical significance of the results.

## Data availability

All data generated and analyzed in this study are included in the article and its Supplementary Information, and are also available from the authors upon request.

## Acknowledgments

This work was supported by a Pinnacle Research Award from American Association for the Study of Liver Disease (J.-Y.S.); Gilead Science Research Scholar Award (J.-Y. S.); Columbia University Digestive and Liver Diseases Research Center Pilot Grant (J.-Y.S.); and the National Institutes of Health [grant numbers R01DK118480 (W.T.D., H.N.G., H.J.W.), R35HL135833 (H.N.G.)]. This research was supported by the Columbia University Digestive and Liver Disease Research Center grant 1P30DK132710. Some of the image processing and analysis was performed in the Confocal and Specialized Microscopy Shared Resource of the Herbert Irving Comprehensive Cancer Center at Columbia University, which is supported by the National Institutes of Health [grant number P30CA013696]. The content is solely the responsibility of the authors and does not necessarily represent the official views of the National Institutes of Health.

## Author contributions

W.T.D, H.N.G., H.J.W. and J.-Y. S. conceived of the project. A.H.-O., W.T.D, H.N.G., H.J.W. and J.-Y.S. designed experiments. W.T.D. generated mouse lines. J.-Y.S designed and generated the viral construct. A.H.-O., Y.P.Z., J.W. M., C.Ö, M. J. L. and J.-Y.S. performed experiments. A.H.-O., Y.P.Z., J.W.M., A.S., H.N.G., H.J.W. and J.-Y.S. analyzed data. W.T.D, H.N.G., H.J.W. and J.-Y.S obtained funding. A.H.-O., Y.P.Z., W.T.D, H.N.G., H.J.W. and J.-Y.S wrote the manuscript. All authors reviewed and contributed to the final manuscript.

## Additional information

Supplementary information The online version contains supplementary material available at xxxx

**Correspondence** and requests for materials should be addressed to Howard J. Worman, Henry N. Ginsberg and Ji-Yeon Shin.

