## supplemental materials for "Functional interaction of torsinA and its activators in liver lipid metabolism"

[1]Department of Medicine, Vagelos College of Physicians and Surgeons, Columbia University, New York, NY, USA. [2]Columbia Center for Human Development, Columbia University Irving Medical Center, New York, NY, USA. [3]Department of Pathology and Cell Biology, Vagelos College of Physicians and Surgeons, Columbia University, New York, NY, USA. [4]Department of Biostatistics, Mailman School of Public Health, Columbia University, New York, NY, USA. [5]Peter O'Donnell Jr. Brain Institute, University of Texas Southwestern Medical Center, Dallas, TX, USA. [6]Department of Neurology, University of Texas Southwestern Medical Center, Dallas, TX, USA. [7]Department of Neuroscience, University of Texas Southwestern Medical Center, Dallas, TX, USA. [8]These authors contributed equally: Antonio Hernandez-Ono, Yi Peng Zhao. [9]Correspondence: Howard J. Worman, Henry N. Ginsberg, Ji-Yeon Shin.

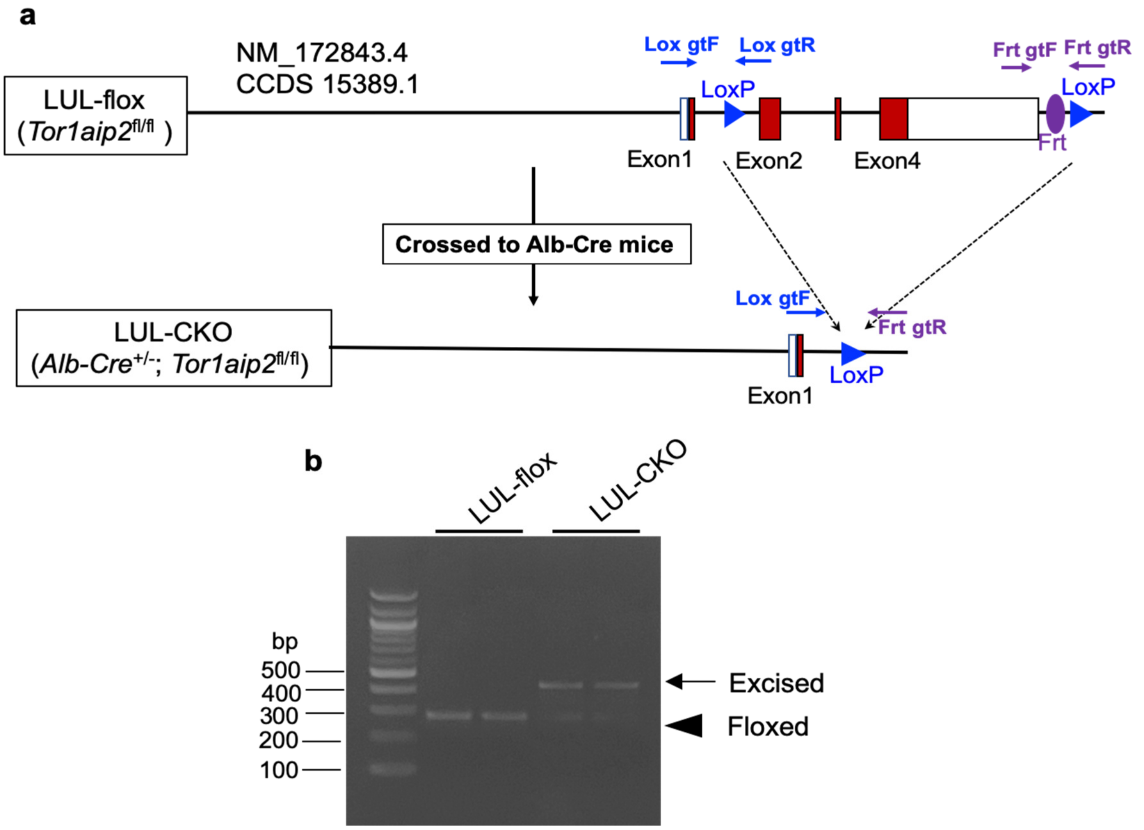


**Supplementary Fig. 1. Generation of LUL-CKO mice. a** To generate LUL-flox mice, the *Tor1aip2* locus (NM_172843.4 or CCDS 15389.1) encoding LULL1 was targeted and engineered to have two loxP sequences (blue triangle) flanking exon 2 to exon 4 (exons shown in red) and a Frt sequence (purple circle) after the end of 3' UTR region of the gene. Blue arrows indicate primers Lox gtF and Lox gtR to detect the first LoxP site located in an intron between exon 1 and exon 2. Purple arrows indicate primers Frt gtF and Frt gtR to detect the Frt sequence. The sequences of corresponding primers are listed in the Supplementary Table 1. To generate LUL-CKO mice with depletion of LULL1 from hepatocytes, we crossed the LUL-flox mice to Alb-Cre transgenic mice to excise the LoxP flanking region. **b** SYBR Green-stained agarose gel showing PCR products demonstrating Cre-mediated DNA excision in mouse livers. Genomic DNA was extracted from livers of LUL-flox and LUL-CKO mice. PCR was performed using primers (Lox gtF, Frt gtR and Frt gtR) that detect the floxed (253 bp, arrow head) and excised (420 bp, arrow) alleles. Both floxed and excised alleles were detected from LUL-CKO samples due to the presences of other non-hepatocyte cells. Molecular mass standards (DNA ladder) are shown in the leftmost lane of the gel.

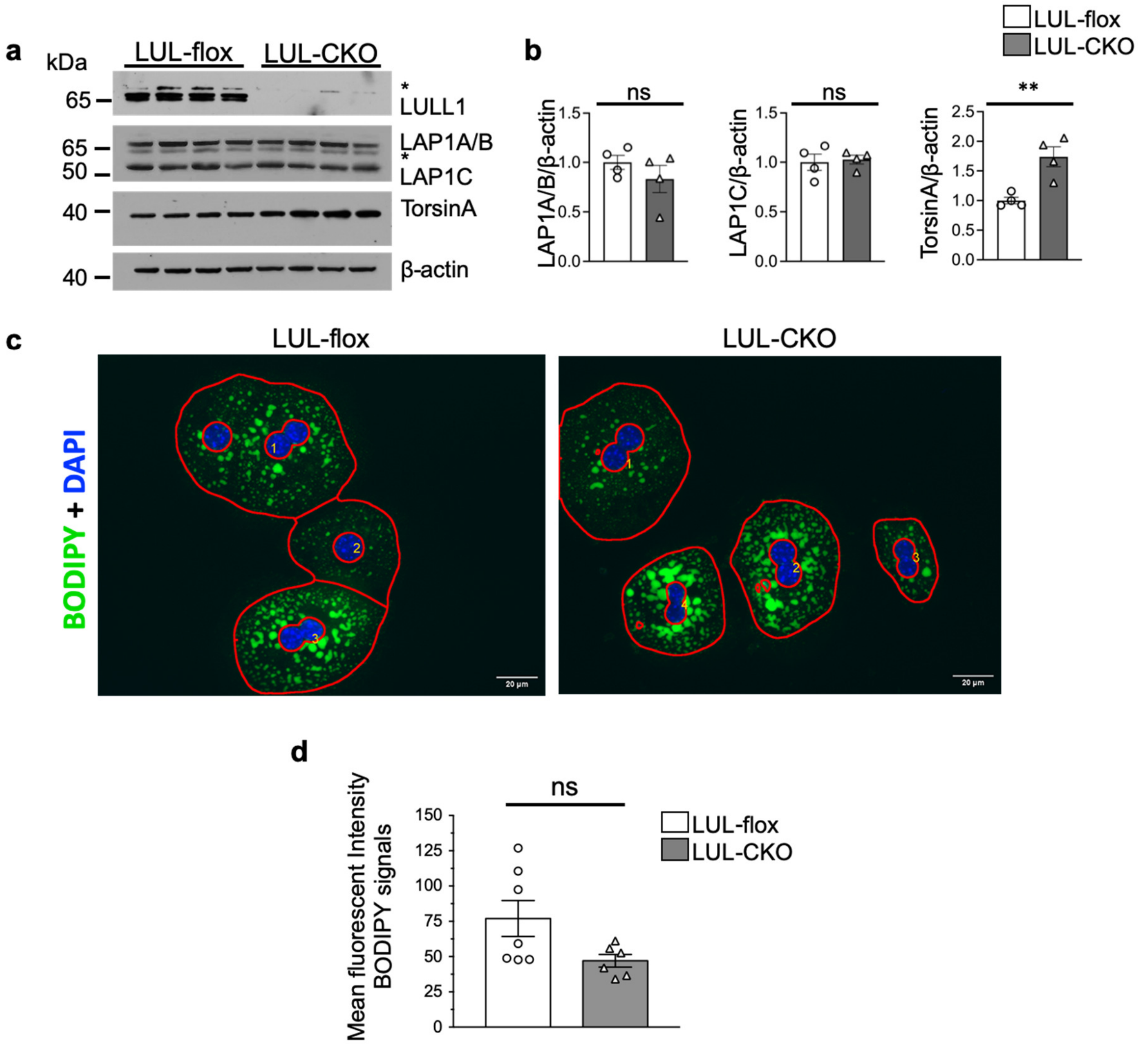


**Supplementary Fig. 2. Modest increase of torsinA expression in livers from LUL-CKO mice and normal lipid droplet distribution in hepatocytes isolated from LUL-CKO mice. a** Immunoblots of liver lysates from LUL-flox and LUL-CKO mice at 4-6 months of age. Blots were probed with antibodies against LULL1, LAP1, torsin, and β-actin. The anti-LULL1 antibody detects two closely migrating bands (see reference 8 in main article). Mice express three LAP1 isoforms — LAP1A, LAP1B, and LAP1C — with LAP1A and LAP1B varying by only 19 amino acids. *Indicates non-specific bands often detected in whole liver lysates using polyclonal antibodies against LAP1 or LULL1. Each lane is a lysate from an individual mouse. **b** Relative ratios of band densities of LAP1A/B, LAP1C, or torsinA to β-actin blots shown in panel (n=4 mice per group). Band densities were measured from immunoblots using protein extracts from four mice per group. Graphs are means ± SEM, Individual circle and triangle is the value from each lane of the immunoblot shown in panel a. ns = not significant, $**p < 0.01$ by Student's t-test. **c** Confocal micrographs of primary hepatocytes from LUL-flox and LUL-CKO mice stained with BODIPY and DAPI. The red lines marked cell boundary and excluded the nuclear region for measurement of cytosolic fluorescent intensity. Numbers with white color indicate counting of the cells during automated scoring (see Methods). **d** Mean fluorescence intensity plots of BODIPY signals calculated from 7 different images of total 48 hepatocytes from LUL-flox and 6 different images of total 35 hepatocytes from LUL-CKO mice. Data are means ± SEM. Circles and squares indicate mean intensity values of each images. ns = not significant by Student's t-test.

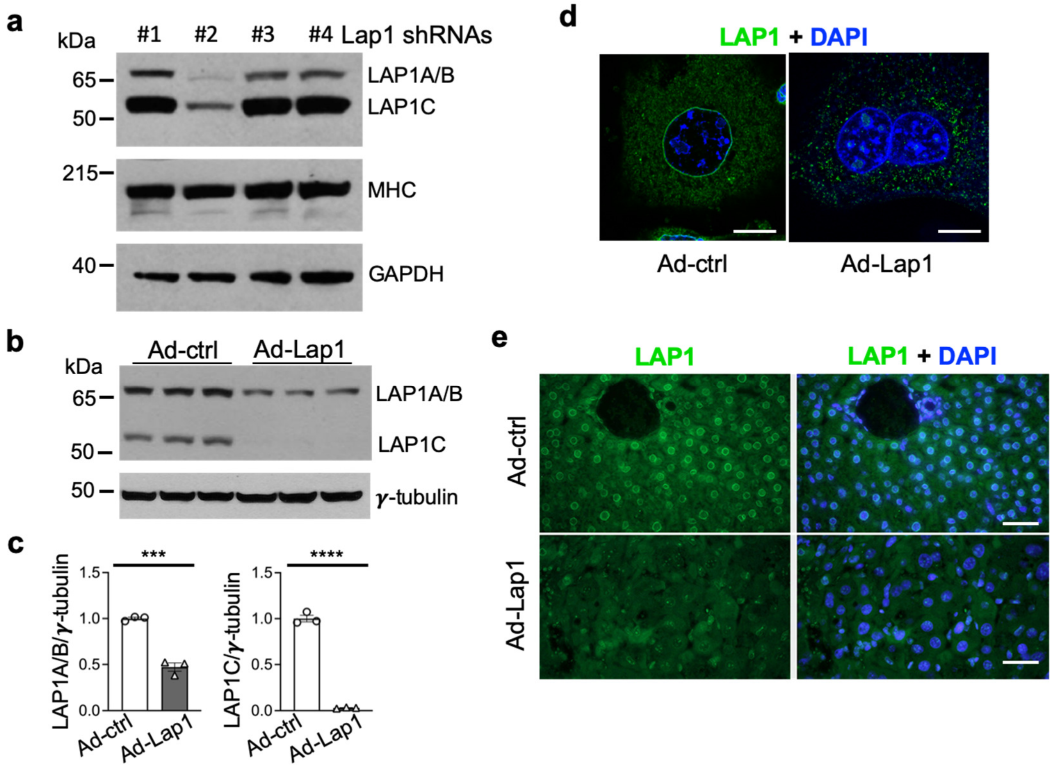


**Supplementary Fig. 3. Validation of adenoviral vectors expressing shRNA to deplete LAP1 from hepatocytes. a** Immunoblots of C2C12 cell lysates collected 48 hours after transduction of retroviral constructs expressing four different shRNAs that target LAP1. Blots were probed with antibodies against LAP1, myosin heavy chain (MHC), and GAPDH. Lap1 shRNA #2 was selected as it gave the greatest LAP1 depletion. **b** Immunoblots of primary hepatocyte lysates probed with anti-LAP1 and anti β-actin antibodies. Hepatocytes isolated from a wild type mouse were transduced with adenoviral vectors that expressed either control scrambled shRNA (Ad-ctrl) or shRNA that targets LAP1 (Ad-Lap1) in a dose of multiplicity of infection 50 at the time of plating and collected 48 hours after transduction. **c** Relative ratios of band densities of LAP1A/B and LAP1C to γ-tubulin in immunoblots shown in panel b. Values are means ± SEM with each circle or square representing the value from each lane of the immunoblot shown in panel b. ****P* <0.001 , *****P* <0.0001 by Student's t-test. **d** Confocal immunofluorescence micrographs of hepatocytes isolated from a wild type mouse administered the indicated adenovirus construct at a dose of 2 x $10^9$ plaque-forming- units/kg. Hepatocytes were labeled anti-LAP1 antibody (green) and DAPI (blue). Scale bars: 10 µm. **e** Immunofluorescence micrographs of liver sections stained anti-LAP1 antibody (green) and DAPI (blue). Five days post injection of wild type mice with Ad-ctrl or Ad-Lap1 shRNA, liver tissues were collected and fixed. Scale bars: 50 µm.

**Supplementary Table 1**. Sequences of primers used for genotyping of LUL-flox and LUL-CKO mice

| **Primer (see Supplementary Fig. 1)** | **Sequence** |
|---|---|
| Lox gtF | acgggaaacaaaaggaggtt |
| Lox gtR | gatggctcagcggttaagag |
| Frt gtF | aatgggtgtgcggttgtta |
| Frt gtR | tggtcaagagagagccaggt |

**Supplementary Table 2.** Quantification of mRNAs for LAP1B/C, LULL1 and HPRT in mouse liver*

| Encoded Protein | Mean CT | SEM |
|---|---|---|
| LAP1B/C | 22.83 | 0.07 |
| LULL1 | 23.25 | 0.06 |
| HPRT | 25.63 | 0.08 |

*Total RNAs were used for quantitative real-time PCR. Mean cycle threshold (CT) values and SEM were calculated from values from 4 different mice per group. The primers for LAP1 detect mRNAs for isoforms LAP1B and LAP1C. The sequences of primers used are provided in the Supplementary Table 3.

**Supplementary Table 3.** Primers used for quantitative real-time PCR to detect mRNAs encoding LAP1B/C, LULL1 and HPRT

| **Protein** | **Sequence** |
|---|---|
| LAP1B/C forward primer | tgcagacccccattaagaag |
| LAP1B/C reverse primer | ggatctggccctgatagtga |
| LULL1 forward primer | cagcagtggttccctagtttag |
| LULL1 reverse primer | ggcagtagtcagccctgttaat |
| HPRT forward primer | agttgagagatcatctccac |
| HPRT reverse primer | ttgctgacctgctggatttac |

### Supplemental Materials and Methods

#### Quantitative real-time qPCR

Total RNA was extracted from dissected livers from mice 4-6 months of age using the RNeasy Isolation Kit (Qiagen). Quality and concentrations of RNA were measured using a Nanodrop spectrophotometer (Thermo Fisher Scientific). The cDNA was synthesized using Superscript First Strand Synthesis System instructions (Thermo Fisher Scientific). For each replicate in each experiment, RNAs from livers of different animals were used (n = 4 per group). The sequences of qPCR primers were either designed using Primer3 software or used with validated primers from the PrimerBank [1]. PCR was performed on a MX3005p qPCR System (Stratagene) using Brilliant III Ultra Fast SYBR Green qPCR Master Mix (Agilent Technologies, 600883). Relative levels of mRNA expression were calculated using the ΔΔCT method [2]. Individual expression values were normalized by comparison to mRNA for HRPT.

#### Supplementary Materials References

1. Wang, X., Spandidos, A., Wang, H. & Seed, B. Primerbank: A pcr primer database for quantitative gene expression analysis, 2012 update. *Nucleic Acids Res.* **40,** D1144-D1149 (2012).

2. Ponchel, F., *et al.* Real-time pcr based on sybr-green i fluorescence: An alternative to the taqman assay for a relative quantification of gene rearrangements, gene amplifications and micro gene deletions. *BMC Biotechnol.* **3,** 18 (2003).
